# Suppression of CD4 T cells by Tregs dictates metabolic remodeling of CD8 T cells during effector to memory transition

**DOI:** 10.64898/2026.02.01.703100

**Authors:** Adithya K. Vegaraju, Vandana Kalia, Surojit Sarkar

**Author notes:** Corresponding authors: Surojit Sarkar and Vandana Kalia.

## Abstract

Metabolic transitions between naïve, effector and memory T cell states are largely orchestrated by TCR, costimulatory and cytokine signals along with nutrient availability in the immune microenvironment. Treg cells have been shown to play a critical role in the effector-to-memory (E-M) transition of virus-specific CD8 T cells through regulation of proliferation and cytotoxic functional programs. However, the precise Treg-dependent metabolic changes that occur in the microniches and, the underlying molecular and cellular mediators of E-M transition remain undefined. Here we show that Treg cells promote the metabolic remodeling of memory-precursor effector CD8 T cells (MPEC) from aerobic glycolysis to fatty acid oxidation as they enter quiescence after antigen clearance. Our data implicate the anatomic microniche of the splenic white pulp as a site for Treg-MPEC interactions. We further show that optimal E-M metabolic transition requires regulation of effector CD4 T cells and inflammatory myeloid cells through inhibitory CTLA4 signals from Treg cells. Moreover, antagonism of inflammatory cytokine interferon-γ (IFN-γ) signals partially rescues the memory defects associated with absence of Treg cells. Together, these findings support a metabolic triad model of memory CD8 T cell differentiation where Treg-dependent regulation of inflammation from effector CD4 T cells promotes the transition of CD8 T cells from cytotoxic effector to quiescent memory metabolic programs. These studies define novel molecular targets that may be exploited to manipulate metabolism, migration and memory function during vaccination.

## INTRODUCTION

Long-lived, polyfunctional memory CD8 T cells that form after an acute infection are critical for rapid and effective pathogen control in case of secondary encounters. Mounting evidence from genetic marking, multiomics and epigenetic studies support a linear developmental path of memory cells from effector CTLs (1–11). After resolution of infection, the memory precursor effector cells (MPECs) slowly convert from a functional effector state to a quiescent memory state by downregulating their effector and proliferative programs and progressively acquiring hallmark memory properties of antigen-independent homeostatic self-renewal and recall expansion (12). This transition from a state of active cytotoxicity to quiescence was thought to occur passively on “autopilot” in the absence of antigen (13–15) owing to lack of continued TCR stimulation. Nonetheless, it is being increasingly recognized that cell-intrinsic regulatory programs such as inhibitory PD-1 signals (16, 17) as well as cell-extrinsic immunosuppressive Treg cells exert additional critical roles in promoting the development and maintenance of quiescence in memory-fated CD8 T cells by suppressing the effector and proliferative programs in developing memory cells (18, 19).

During priming and effector CTL expansion, Treg cells promote the development of long-lived memory CD8 T cells by restraining excessive inflammatory (such as IL-12, IFN-I) and costimulatory signaling (such as CD28, PD-1), and by restricting IL-2 availability to limit terminal effector differentiation (20–26). Treg-derived TGF-β and IL-10 further restrain effector CTL differentiation and promote memory and tissue-resident fates during the expansion phase. However, our understanding of the molecular mechanisms by which Treg cells promote the effector-to-memory transition remains limited. Treg-derived TGF-β has been implicated in the process (27). Moreover, Treg cell-derived IL-10 has been shown to calm inflammation and the activation state of dendritic cells during the effector-to-memory transition phase, thereby promoting memory development (19). Likewise, the inhibitory receptor CTLA4 (expressed at high levels on Tregs) has been also shown to calm inflammation, thereby supporting the downregulation of proliferative and effector programs in MPECs (18).

In this study we investigated how anatomic micro niches and interactions with other immune cells (such as CD4 T cell help) impact Treg-dependent metabolic remodeling of memory-fated effector CD8 T cells as they enter quiescence after antigen clearance. In the murine model of acute LCMV infection, intravascular staining showed a preferential enrichment of MPECs and Treg cells in the splenic T cell zones (white pulp) during the memory transition phase. Using genetically targeted Foxp3^DTR^ mice to specifically ablate Treg cells expressing the human diphtheria toxin receptor (DTR), our studies show that MPEC localization in the splenic white pulp, their metabolic state and protective functional outcomes are dysregulated in the absence of Treg cells. Dysregulation of E-M CD8 T cell metabolism in the absence of Treg cells was in part related to increased CD4 and myeloid T cell activation and induction of pro-inflammatory IFN-γ, which was reversed by soluble CTLA4. Collectively, these studies support the four-cell model of E-M transition, where CTLA4 on Treg cells blocks CD28 signals to CD4 and CD8 T cells by engaging the CD80/86 on antigen presenting cells to maintain homeostasis and promote quiescence. These findings may guide novel strategies for accelerating memory differentiation following immunization.

## MATERIALS AND METHODS

### Animals

Four-week-old C57BL/6 mice were purchased from the Jackson Laboratory (Sacramento, CA, USA). Thy1.1+ and Ly5.1+ H-2D^b^GP33-specific TCR-transgenic P14 mice, fully backcrossed onto the C57BL/6 background, were maintained in our colony. FoxP3^DTR^ mice were purchased from Jax and bred in house as well. All mice used in experiments were 6-8 weeks old and sex-matched within each experiment. Mice were housed in the vivarium at Seattle Children’s Research Institute, and all experiments were conducted in accordance with Institutional Animal Care and Use Committee (IACUC) guidelines.

### Adoptive Transfers

To generate P14 chimeric mice, naïve C57Bl/6 or FoxP3^DTR^ mice were adoptively transferred with 1×10^5^ WT D^b^GP33-specific P14 CD8 T cells 12-16 hrs. prior to infection with LCMV_Arm_.

### Virus and Infection

The Armstrong strain of LCMV (LCMV_Arm_) was propagated in-house as described previously(28). Mice were infected with 2×10^5^ plaque-forming units of LCMV intraperitoneally to initiate an acute infection as previously(29).

### Metabolic assays

Mitochondrial assays were performed with samples after isolation using tetramethylrhodamine ethyl ester (TMRE) and Mitotracker (Invitrogen) per manufacturer’s instructions. Fluorescent glucose analog N-(7-Nitrobenz-2-oxa-1,3-diazol-4-yl)Amino)-2-Deoxyglucose (2-NBDG) (Invitrogen) was used for glucose flux analyses per manufacturer’s instructions. Mitosox was used for assessing mitochondrial superoxides per manufacturer’s protocol.

Briefly, for 2-NBDG labeling, 1×10^6^ cells were placed in 96-well plates and washed with glucose-free media (1800 RPM, 2 minutes). Then, 50μL of 2-NBDG diluted in glucose-free media was added to the cells and incubated for 30 minutes at 37°C. Following this step, cells were labeled with fluorescent antibodies as previously described, fixed in 4% paraformaldehyde, and analyzed by flow cytometry.

For MitoSox labeling, 1×10^6^ cells were placed in 96-well plates and were washed with 10% RPMI medium. Then, 50μL of MitoSox dye diluted in 10% RPMI medium was added to the cells and incubated for 30 minutes at 37°C. Following this step, cells were labeled with fluorescent antibodies as previously described and fixed in 4% paraformaldehyde prior to FACS analysis.

For Mitotracker and TMRE labeling, 1×10^6^ cells were placed in 96-well plates and were washed with 10% RPMI medium. Then, 50μL of a mixture containing fluorescent antibodies, Mitotracker and TMRE dyes diluted in 10% RPMI was added to the cells and incubated for 30 minutes at 37°C. The cells were washed once with PBS and analyzed by flow cytometry immediately.

### Depletion of Regulatory T-cells

FoxP3DTR mice and Diphtheria Toxin (DT) was used to deplete regulatory T-cells in mice. Injection of 20ng DT/g every three days starting day 8 post-infection was used to deplete regulatory T cells in mice after antigen clearance. DT was purchased from Sigma Aldrich.

### CD4 T cell Depletion

CD4 T cell depletion was achieved by injecting 500μg of αCD4 (GK1.5) CD4 depleting antibody on days 8 and 9 post-infection. GK1.5 was purchased from BioXcell.

### CTLA4-Ig Treatment

CTLA4-Ig was kindly provided by Dr. Steven G. Nadler. 500μg was injected into mice every three days starting at day 8 post-infection.

### Intravascular Cell Staining

In order to differentiate between CD8+ T cells in red pulp and white pulp of the spleen, the mice were injected with fluorochrome-conjugated anti-CD8β (clone 53-6.7) intravenously and euthanized 4 minutes later. Lymphocytes were isolated from indicated tissues, followed by staining with anti-CD8α antibody. Cells positive for CD8α and CD8β were circulatory in the red pulp of the spleen and the CD8α+ CD8β-population was considered to be in the white pulp.

### Flow cytometry and cell sorting

All antibodies were purchased from Biolegend (San Diego, CA, USA), except granzyme B, which was purchased from Invitrogen. MHC class I tetramers were made as described in NIH tetramer core facility protocol. Cells were stained for surface proteins in 1xPBS containing 1% FBS and 0.05% sodium azide. Intracellular proteins and cytokines were stained using BD intracellular flow kit (BD biosciences). For analysis of intracellular cytokines, 2 × 10^6^ lymphocytes were stimulated with 0.2 μg/mL GP_33-41_ peptide in the presence of Brefeldin A (BFA) for 5 h, followed by surface staining for CD8, Ly5.1, Thy1.1, and Thy1.2, and intracellular staining for IFN-γ, TNF-α, or IL-2. Single cell suspensions of spleen (SPL), inguinal lymph nodes (iLN), lungs (LNG), livers (LVR) or PBMCs were prepared and direct *ex vivo* staining was carried out as reported previously(17). LSRII Fortessa (BD Biosciences, San Jose, CA) was used for flow cytometry and analysis was done using FlowJo software (Treestar).

For cell sorting CD8 T cells were enriched from total splenocytes using the EasySep Mouse CD8+ T cell Isolation Kit (Biolegend), at day 12 post infection. The enriched CD8 T cells were then FACS-sorted into CD8β+ (red pulp) and CD8β- (white pulp) cells, following gating of Thy1.2- CD44+ CD127^Hi^ P14 population. Cells were sorted on JAZZ cell sorter (BD Bioscience) using a 70-micron nozzle.

### Statistical Analysis

Statistical analysis was performed using Student’s t-test for two experimental groups and 1-way ANOVA for multiple experimental groups. A paired t-test was used in any comparison between red pulp and white pulp values, and unpaired t-test was used for comparison between two groups. P values of statistical significance are depicted by an asterisk: *(P ≤ 0.05), ** (P ≤ 0.01),*** (P ≤ 0.001). (P > 0.05) was considered not significant.

## RESULTS

### Treg cells and MPECs are preferentially enriched in the splenic white pulp T cell zones

Terminal effector and memory CD8 T cells localize to different anatomic locations in the spleen – with effectors preferentially localizing in the red pulp and memory cells enriched in the white pulp T cell zones (30). Moreover, MPECs localizing within the splenic white pulp have been shown to differentiate into higher quality memory cells than those from the red pulp (27). However, location-specific factors that may drive differential memory differentiation programs remain unknown. Proximity to regulatory T-cells has been hypothesized as a possible reason for this differential outcome in memory quality. Hence, we first sought to determine the anatomic localization of splenic Tregs and MPECs during the E-M transition phase following an acute LCMV infection using intravascular staining. We observed preferential enrichment of Treg cells (Fig 1A) and CD127^Hi^ KLRG-1^Lo^ MPEC cells (Fig 1B) in the splenic white pulp (∼60%) compared to red pulp (∼30%) at day 12 post-infection – a time when virus is fully cleared and MPECs begin to acquire memory properties (12, 18, 31). In contrast, CD127^Lo^ KLRG-1^Hi^ shortlived effector cells (SLECs) preferentially localized in the splenic red pulp at day 12 (∼40% in red pulp compared to 10% in white pulp) (Fig 1B). Similar results were noted at days 14 and 22 post-infection as well (Fig S1). Notably, no localization differences were observed for MPECs or SLECs at the peak of clonal expansion (day 8 post-infection) (Fig S1), with over 70% terminal effectors and ∼5% MPECs localized similarly in both red and white pulp areas. Importantly, consistent with previous studies, we observed better recall expansion potential of memory-fated cells isolated from splenic white pulp compared to red pulp (Fig 1C). These data show that differences in anatomical localization of MPECs and SLECs arise in the early E-M transition phase, with MPECs preferentially enriched in the white pulp area along with Treg cells, and SLECs preferentially localized in the red pulp.

**Figure 1.**
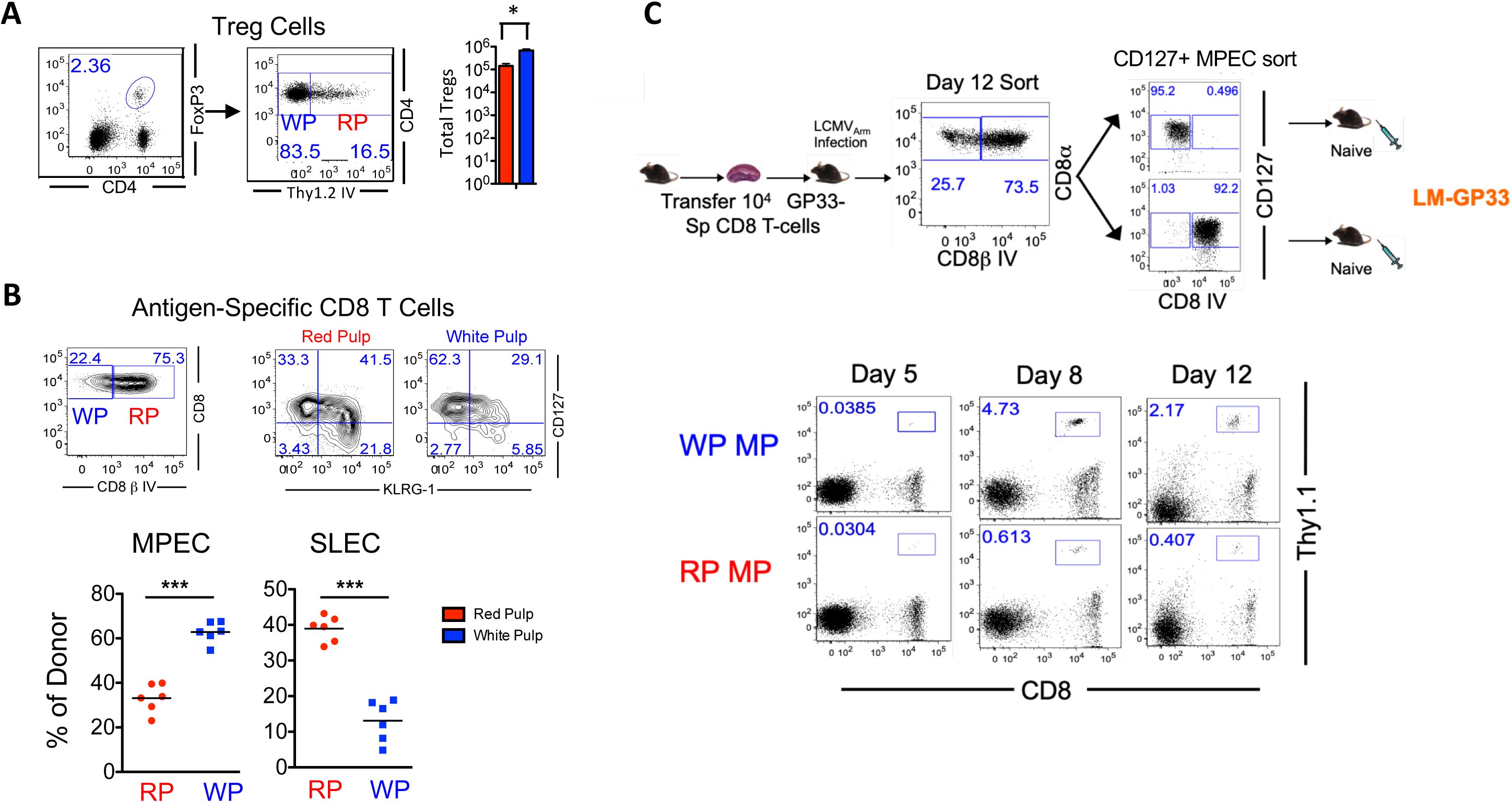
Treg cells and MPECs are preferentially enriched in the splenic white pulp areas. A. Anatomic localization of Tregs in spleen on day 12 post-infection with LCMV_Arm_. 1×10^5^ D^b^GP33-specific P14 T cells were adoptively transferred into C57BL/6 mice which were subsequently infected with LCMV_Arm_. Intravascular staining with fluorescent Thy1.2 antibody was performed to differentiate the red pulp and white pulp cells of the spleen. Raw FACS plots and composite bar graphs depict the red pulp and white pulp distribution of CD4+FoxP3+ Treg cells. B. Frequency of D^b^GP33-specific CD127^hi^KLRG-1^lo^ MPECs and CD127^lo^KLRG-1^hi^ SLECs in splenic anatomic compartments at day 14 post-infection. 1×10^5^ D^b^GP33-specific P14 CD8 T cells were adoptively transferred into C57BL/6 mice which were subsequently infected with LCMV_Arm_. Intravascular staining with fluorescent CD8β antibody was performed to differentiate P14 CD8 T cells in the red pulp and white pulp splenic compartments. Cell surface markers CD127 and KLRG-1 were used to assess proportions of MPEC and SLEC subsets as presented in the representative FACS plots. C. CD127^Hi^ memory-fated P14 CD8 T cells were isolated from splenic red and white pulp using intravascular staining, and adoptively transferred in equal numbers (∼5×10^3^) into congenically mismatched naïve B6 mice. The mice were rechallenged with recombinant *Listeria monocytogenes* expressing LCMV GP33 epitope (LM-GP33) and donor cell expansion was assessed longitudinally by flow cytometry in PBMC samples at days 5, 8 and 12 post rechallenge. Data are representative of 2-5 experiments with 3-5 mice per group. Statistical significance is indicated by the following: *p ≤ 0.05, **p ≤ 0.01, ***p ≤ 0.001. p ≥ 0.05 was considered ns.

### Treg cells promote the maintenance of MPECs in splenic microniches

We next queried whether the preferential localization of MPECs and their memory potential in the white pulp was dependent on the presence of Treg cells. To test this hypothesis, we leveraged FoxP3^DTR^ mice to specifically deplete Treg cells during E-M phase using diphtheria toxin. Treg cells were specifically ablated using diphtheria toxin treatment starting at day 8 after acute LCMV infection (Fig S2A), which marks the beginning of the E-M phase. As shown in Fig 2A and 2B, Treg cell depletion led to reduced proportions of MPECs and increased proportions of SLECs in both white pulp and red pulp regions of the spleen. However, Treg cells prove necessary for the maintenance of total numbers of MPECs in all splenic compartments (Fig 1B). There were significantly fewer total MPECs in the absence of Tregs in both white and red pulp zones. SLECs, on the other hand, maintained their numbers similarly in the presence or absence of Treg cells in both the red and white pulp areas. Loss of Tregs led to a specific loss of MPECs, but not SLECs in both red and white pulp areas at all time points analyzed during the E-M transition phase (Fig S2B).

**Figure 2.**
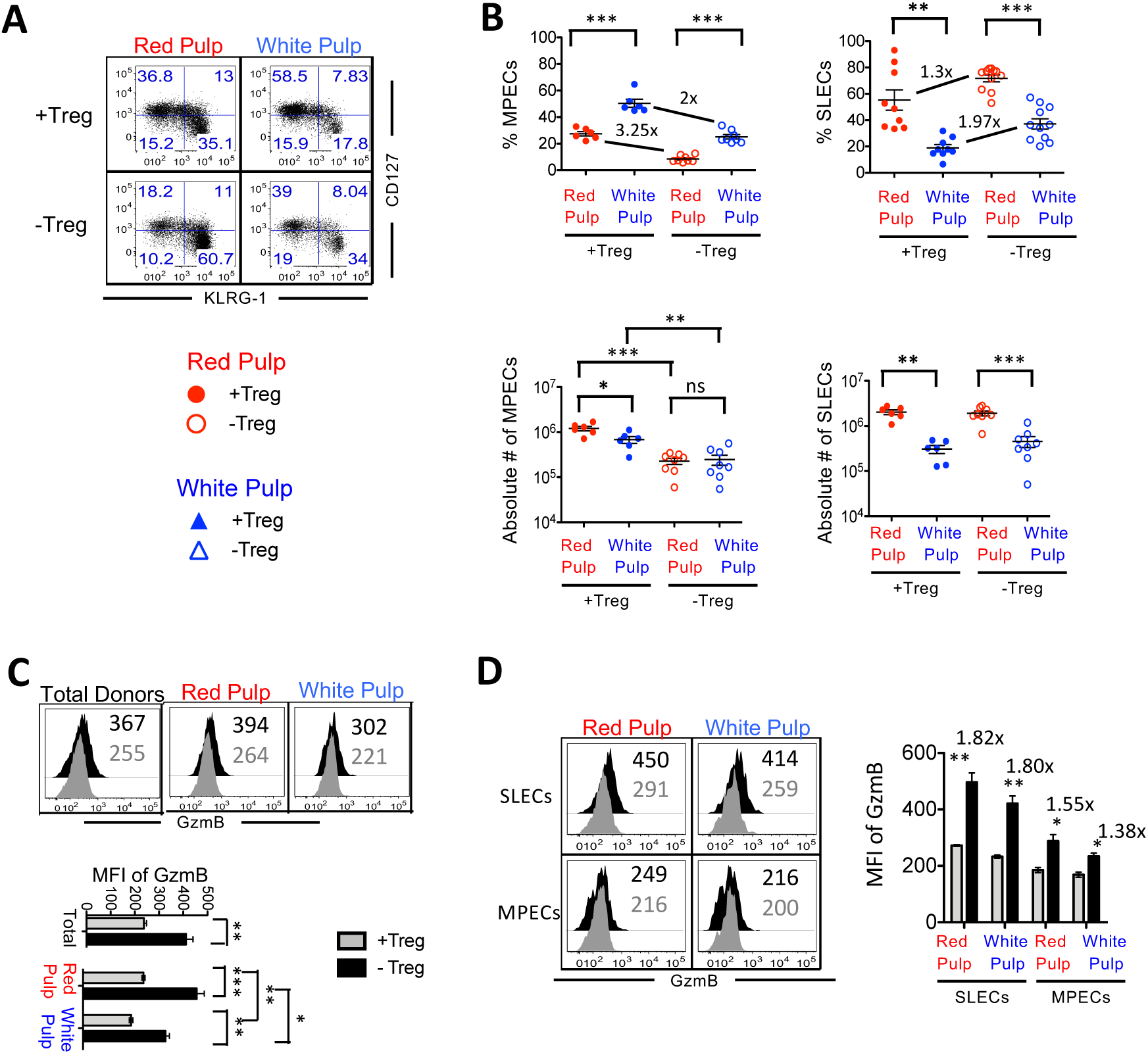
Maintenance of MPECs is critically dependent on Treg cells. 1×10^5^ D^b^GP33-specific P14 T cells were adoptively transferred to C57BL/6 (+Treg) and Foxp3^DTR^ (-Treg) mice which were subsequently infected with LCMV_Arm_. FoxP3^DTR^ mice were treated with Diphtheria Toxin (DT) every three days starting day 8 to deplete Treg cells. Intravascular staining with fluorescent CD8β antibody was performed to differentiate P14 CD8 T cells in the red pulp and white pulp splenic compartments. A. Percentage of donor CD127^hi^KLRG-1^lo^ MPECs in splenic anatomic compartments on Day 14 p.i. with LCMV_Arm_ under Treg sufficient and deficient conditions. The flow plots show MPEC and SLEC distribution of the red pulp and white pulp donors in the presence and absence of Tregs on Day 14. B. The scatter plot depicts the change in MPEC and SLEC enrichment within each anatomic compartment as well as absolute numbers of donor MPECs and SLECs at day 14 p.i. upon Treg cell depletion. C. Granzyme-B (GzmB) expression in CD8+ donors from red pulp and white pulp of spleen on Day 14 p.i. with LCMV_Arm_ under Treg sufficient and deficient conditions. The histogram compares the mean fluorescent intensities (MFI) of GzmB in CD8+ donors between the described experimental groups. The bar graph depicts the mean fluorescence intensity of granzyme-B in CD8+ donors of both experimental groups. D. Granzyme-B (Gzm-B) expression in donor MPECs and SLECs from red pulp and white pulp of spleen on Day 14 p.i. with LCMV_Arm_ under Treg sufficient and deficient conditions. The histogram compares the mean fluorescent intensities (MFI) of GzmB in MPECs and SLECs from red pulp and white pulp of spleen with and without Treg cells in environment. The bar graph depicts the mean fluorescence intensity of granzyme-B of red pulp and white pulp MPECs and SLECs of both experimental groups. Data are representative of 2-5 experiments with 3-5 mice per group. Statistical significance is indicated by the following: *p ≤ 0.05, **p ≤ 0.01, ***p ≤ 0.001. p ≥ 0.05 was considered ns.

To query the impact of Treg cells on the transition of MPECs from an effector to a memory phenotype, we analyzed the expression levels of granzyme B – a canonical effector molecule whose expression is progressively downregulated during E-M transition (12). Total antigen-specific CD8 T cells showed higher granzyme B expression in the absence of Treg cells, in both red and white pulp splenic areas with CD8 T cells expressing modestly higher levels of granzyme B in the red pulp than in the white pulp (Fig 2C). This difference could result from higher proportions of terminal SLECs in the red pulp. Finer dissection of granzyme B expression levels in MPECs and SLECs in the red and white pulp areas showed that both MPECs and SLECs upregulated granzyme B expression similarly upon Treg ablation, albeit MPECs expressed lower overall levels of granzyme B compared to SLECs, and a trend towards higher granzyme B expression was noted in the red pulp compared to white pulp (Fig 2D). Collectively, these data demonstrate that Treg cells aid in curbing the effector program in MPECs and SLECs in both splenic red and white pulp areas, and are critically required for maintaining the MPEC numbers in the splenic red and white pulp areas both. The specific loss of MPECs, but not SLECs, upon Treg ablation may be due to MPEC to SLEC conversion in the absence of Treg cells, as suggested previously (18), and/or due to migration of MPECs out of the spleen into the lymph nodes following Treg loss, as suggested by increased numbers of MPECs with increased granzyme B expression in the inguinal lymph nodes following Treg ablation (Fig S2C and S2D).

### Treg cells promote metabolic quiescence of MPECs and SLECs in splenic microniches

During E-M transition, MPECs undergo a major shift in metabolism, switching from glycolysis-based energy production to fatty acid oxidation, which is the preferred mode of energy generation in memory CD8 T cells. To determine whether Treg cells mediate differential metabolic regulation of MPECs and SLECs in the splenic red pulp and white pulp zones, we assessed glucose uptake and mitochondrial metabolic activity of MPECs and SLECs in both red and white pulp areas at day 14 after infection using intravascular staining. The gating strategy is shown in Fig S3A-C. Treg cells were found to inhibit glucose uptake in total splenic donors (Fig 3A). This effect occurs in both red pulp and white pulp donors (Fig 3B), as well as red pulp and white pulp MPECs and SLECs (Fig 3C). This effect is more pronounced in the splenic microenvironment, albeit a similar trend was noted in the iLNs as well as the stroma and parenchyma of the lungs (Fig S3D).

**Figure 3.**
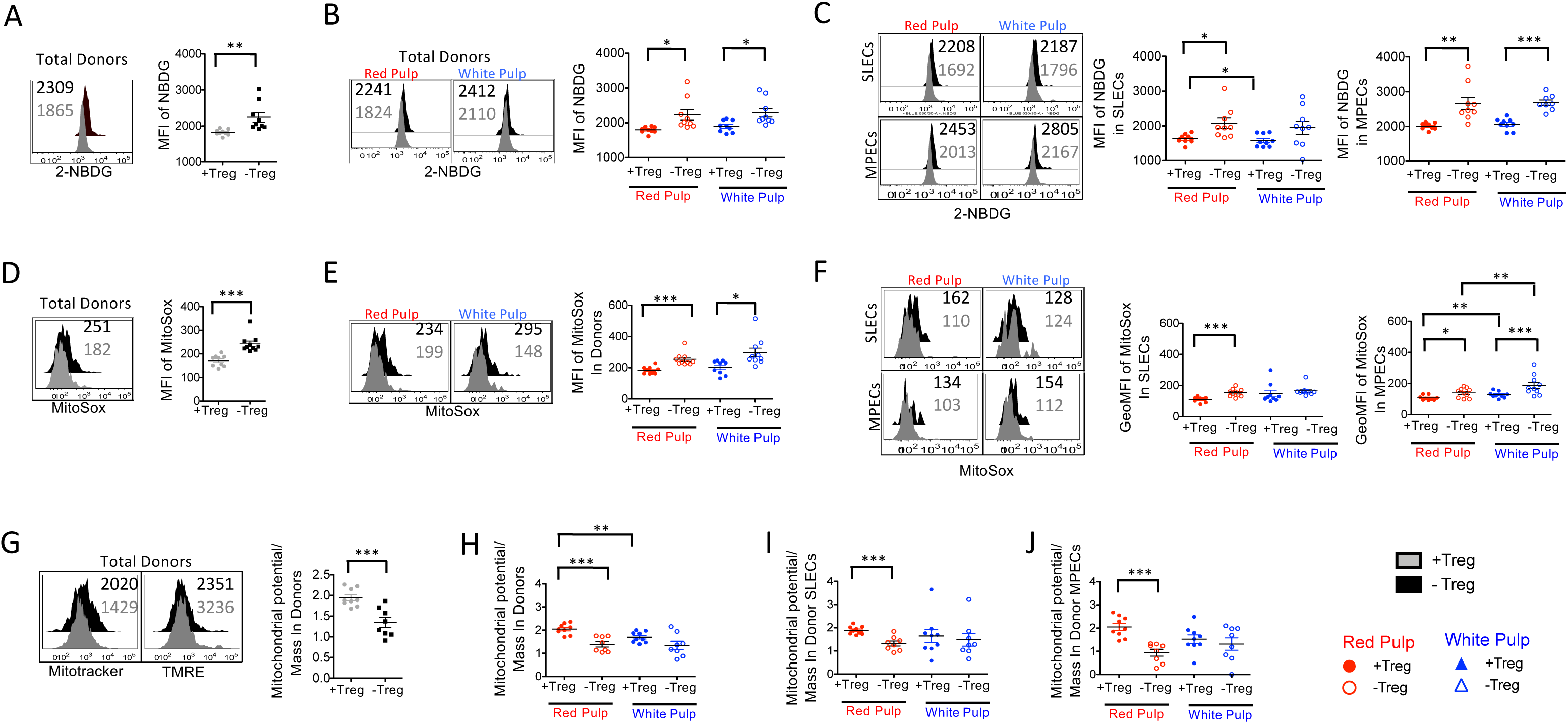
Treg cells promote metabolic quiescence of MPECs and SLECs in splenic microniches. 1×10^5^ D^b^GP33-specific P14 T cells were adoptively transferred into C57BL/6 and FoxP3^DTR^ mice, which were subsequently infected with LCMV_Arm_. FoxP3^DTR^ mice were treated with Diphtheria Toxin (DT) every three days starting day 8 post-infection to deplete Treg cells. Intravascular staining with fluorescent CD8β antibody was performed to differentiate P14 CD8 T cells in the red pulp and white pulp splenic compartments. The mice were harvested on day 14 post-infection and splenic lymphocytes were analyzed with flow cytometry and metabolism assays. A. Glucose uptake in CD8+ T cell donors from Treg sufficient and Treg deficient conditions on day 14 post-infection. FACS sorting was utilized to isolate CD8+ donors from harvested splenic lymphocytes and a glucose uptake assay with 2-NBDG was performed on isolated donors from both experimental groups. The histogram and scatter plot display the MFI of 2-NBDG following glucose uptake assay in donors from Treg sufficient and Treg deficient conditions. B. Oxidative stress in CD8+ T cell donors from Treg sufficient and Treg deficient conditions on day 14 post-infection. Following FACS sorting as described above, oxidative stress was measured with a MitoSox assay on isolated splenic donors from both experimental groups. The histogram and scatter plot display the MFI of MitoSox following the assay in donors from Treg sufficient and Treg deficient conditions. C. Mitochondrial potential and mass in CD8+ T cell donors from Treg sufficient and Treg deficient conditions on day 14 post-infection. Following FACS sorting as described above, mitochondrial mass (Mitotracker) and mitochondrial membrane potential (TMRE) were measured on harvested splenic donors from both experimental groups. The histograms display the MFI values of Mitotracker and TMRE on donors from Treg sufficient and deficient conditions. The scatter plot displays mitochondrial membrane potential per mitochondrial mass unit in donors from Treg sufficient and Treg deficient conditions. D. Glucose uptake in splenic red pulp and white pulp CD8+ T cell donors from Treg sufficient and Treg deficient conditions on day 14 post-infection. FACS sorting was utilized to separate Thy1.2+ (Red Pulp) from Thy 1.2 – (White Pulp) CD8+ donors and a glucose uptake assay with 2-NBDG was performed on the red pulp and white pulp donors from both experimental groups. The histogram and scatter plot display the MFI of 2-NBDG following glucose uptake assay in red pulp and white pulp donors from Treg sufficient and Treg deficient conditions. E. Oxidative stress in splenic red pulp and white pulp CD8+ T cell donors from Treg sufficient and Treg deficient conditions on day 14 post-infection. FACS sorting was utilized as described above to isolate red pulp and white pulp CD8+ donors and a MitoSox assay was performed on the donors from both experimental groups to measure oxidative stress. The histogram and scatter plot display the MFI of MitoSox following the assay in donors from Treg sufficient and Treg deficient conditions. F. Glucose uptake in splenic red pulp and white pulp CD8+ T cell donor MPECs and SLECs from Treg sufficient and Treg deficient conditions on day 14 post-infection. FACS sorting was utilized to separate CD127^hi^KLRG-1^lo^ (MPECs) and CD127^lo^KLRG-1^hi^ (SLECs) CD8 + T cell donors from splenic red pulp and white pulp compartments. A glucose uptake assay with 2-NBDG was performed on the red pulp and white pulp donor MPECs and SLECs from both experimental groups. The histogram and scatter plot display the MFI of 2-NBDG following glucose uptake assay in red pulp and white pulp donor MPECs and SLECs from Treg sufficient and Treg deficient conditions. G. Oxidative stress in splenic red pulp and white pulp CD8+ T cell donor MPECs and SLECs from Treg sufficient and Treg deficient conditions on day 14 post-infection. FACS sorting was utilized as described above to isolate red pulp and white pulp CD8+ donor MPECs and SLECs and a MitoSox assay was performed on the donors from both experimental groups to measure oxidative stress. The histogram and scatter plot display the geoMFI of MitoSox following the assay in donors from Treg sufficient and Treg deficient conditions. H. The scatter plot displays the MFI value of mitochondrial membrane potential per mitochondrial mass unit in donors from Treg sufficient and Treg deficient conditions in red and white pulp areas of spleen. I. The scatter plot displays the MFI value of mitochondrial membrane potential per mito-chondrial mass unit in donor SLECs from Treg sufficient and Treg deficient conditions in red pulp and white pulp. J. The scatter plot displays the MFI value of mitochondrial membrane potential per mito-chondrial mass unit in donor MPECs from Treg sufficient and Treg deficient conditions in red pulp and white pulp. Data are representative of 2-5 experiments with 3-5 mice per group. Statistical significance is indicated by the following: *p ≤ 0.05, **p ≤ 0.01, ***p ≤ 0.001. p ≥ 0.05 was considered ns.

We also assessed mitochondrial metabolic activity using MitoSox (measure of mitochondrial superoxide levels), and mitochondrial potential using TMRE (tetramethylrhodamine ethyl ester – a potentiometric fluorescent dye that measures mitochondrial membrane potential in live cells) and Mitotracker (cell-permeant fluorescent dye that measure mitochondrial mass or content). Excessive or sustained mitochondrial superoxides drive CD8 T cells toward short-lived effector fates and apoptosis, whereas controlled mitochondrial superoxide signaling is associated with memory potential (32, 33). Effector CD8 T cells typically exhibit elevated mitochondrial superoxides, reflecting high electron transport chain flux and reliance on aerobic glycolysis coupled to robust OXPHOS (34, 35). In contrast, memory CD8⁺ T cells display lower basal MitoSOX signal, associated with reliance on fatty acid oxidation and efficient OXPHOS. In addition to regulating glucose uptake, we found that Treg cells also suppressed the levels of mitochondrial superoxide in antigen-specific CD8 T cells (Fig 3D). This effect occurred in both red pulp and white pulp donors (Fig 3E) as well as red pulp and white pulp MPECs (Fig 3F). Notably, the increase in superoxide production in MPECs upon Treg cell ablation was more pronounced in the iLNs and the parenchyma of the lung compared to the spleen (Fig S3E).Similar to the observations on glucose uptake, Treg cell ablation showed insignificant effects on white pulp SLEC mitochondrial superoxide production (Fig 3F).

To further query the role of anatomic location and Tregs on mitochondrial function during E-M, we next assessed the mitochondrial membrane potential of antigen-specific CD8 T cells in splenic red and white pulp, in the presence or absence of Treg cells (Fig 3G-J). Effector CD8⁺ T cells typically exhibit hyperpolarized mitochondria with relatively low mitochondrial mass, whereas memory CD8⁺ T cells are characterized by increased mitochondrial content, fused mitochondrial networks, and lower basal membrane potential, conferring superior respiratory capacity and long-term persistence (36–38). Consistent with this, we observed lower mitochondrial polarization in MPECs in white pulp – memory-fated cells that are associated with optimal memory potential (27, 30) (Fig 3J). Location effects were dominant, such that total antigen-specific CD8 T cells and SLECs in white pulp also exhibited lower mitochondrial polarization than in red pulp (Fig 3H, 3I). Treg cells were found to significantly increase the mitochondrial potential of antigen-specific CD8 T cells, such that higher mitochondrial polarization in total antigen-specific CD8 T cells was observed in the presence of Treg cells, than in their absence (Fig 3G). Interestingly, this effect was observed only in the red pulp, but not white pulp (Fig 3H), such that ablation of Treg cells led to a significant decrease in the membrane polarization of MPECs and SLECs in the red pulp, with minimal effects on white pulp CD8 T cells (Fig 3H-J, S3F-H).

Collectively, our observations that Treg cells regulate glucose metabolism and mitochon-drial oxidative stress, while promoting optimal mitochondrial polarization and mass lend support to the notion that Treg cells support the metabolic transition of effector cells into quiescent memory cells during the E-M transition phase in a location-dependent manner.

### Treg cell dependent E-M transition of MPECs is dependent on CTLA4 and suppression of CD4 T cells

To explore the mechanism by which Treg cells promote E-M transition of antigen-specific CD8 T cells, we studied the effects of total CD4 T cell depletion with or without Treg ablation. We also tested CTLA4-Ig treatment as a surrogate for Treg immunosuppressive functional molecule in Treg cell-depleted mice. As previously shown, when Treg cells were depleted starting at day 8 post-infection (beginning of E-M phase), there was a sharp decline in splenic MPEC numbers with no effect on absolute numbers of SLECs (18) (Fig 4A, S4A). The decline in MPEC numbers occurred in both red pulp and white pulp areas, albeit the effect was more significant in the red pulp. Notably, CD4 T cell depletion completely reversed the deleterious effects of Treg ablation on E-M transition (Fig 4). In the absence of Treg cells, CD4 T cell depletion led to a restoration of MPEC numbers in both the red and white pulp zones (Fig 4A). CD4 T cell depletion (in the absence of Treg cells) also supported down regulation of effector program in splenic red and white pulp, as assessed by downregulation of granzyme B expression during E-M in total donors (Fig 4B, 4C), MPECs (Fig 4D) and SLECs (Fig S4B). As expected, CTLA4-Ig was able to fully replace immunosuppressive Treg functions required for optimal E-M transition of antigen-specific CD8 T cells, as indicated by restoration of MPEC numbers that were lost in the absence of Treg cells (Fig 4A), as well as effective downregulation of the effector gene granzyme B expression in MPECs (Fig 4B-D). These data support a critical regulatory nexus of Treg cells through CTLA4 and effector CD4 T cells during the E-M transition of CD8 T cells into quiescent memory.

**Figure 4.**
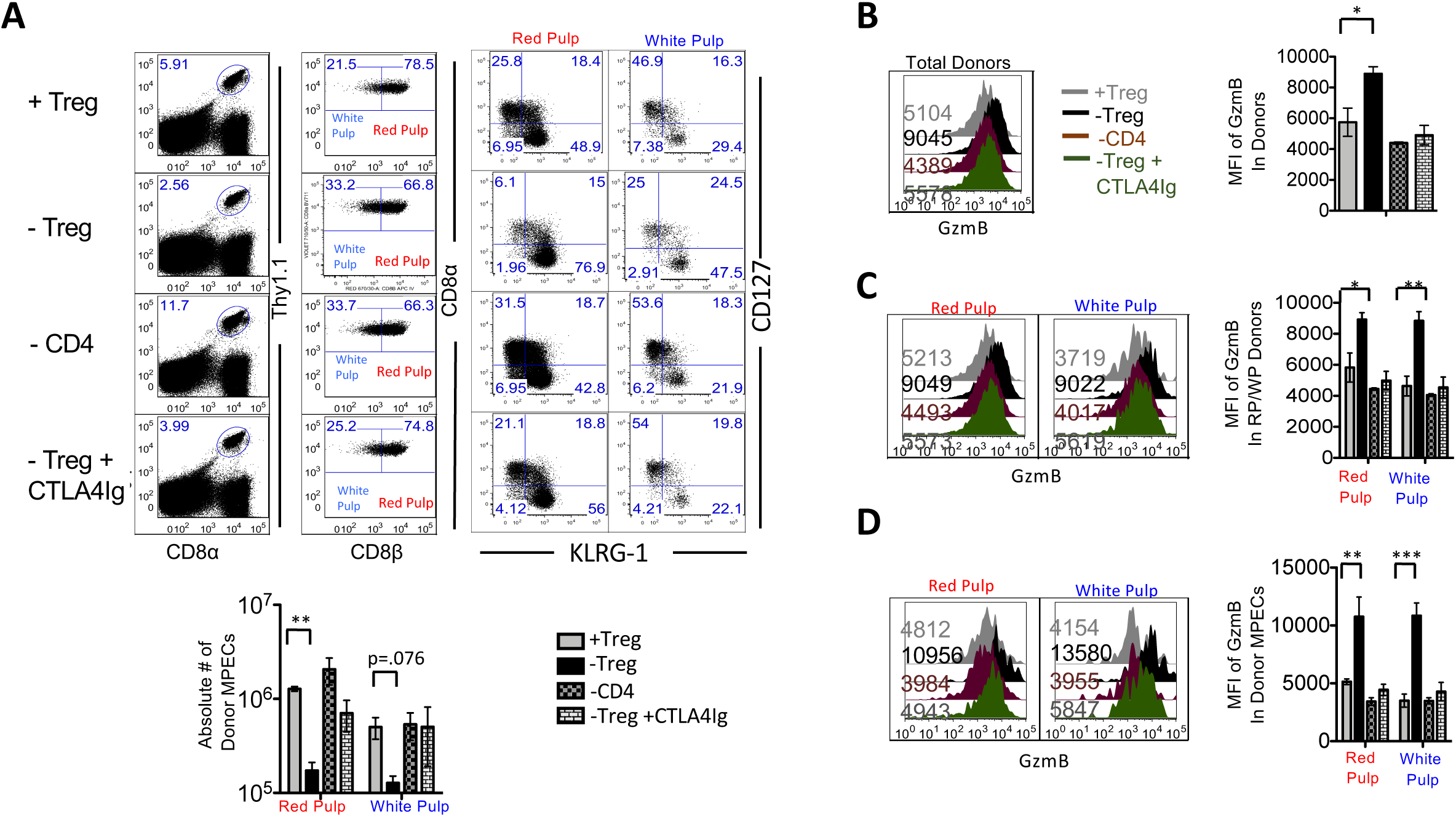
CTLA4 and CD4 T cells exert a critical role in Treg-dependent conversion of effector-to-memory quiescence. 1×10^5^ D^b^GP33-specific P14 T cells were adoptively transferred into Foxp3^DTR^ mice, which were subsequently infected with LCMV_Arm_. Two groups of Foxp3^DTR^ mice were treated with DT on day 8 and day 11 p.i. One of these groups also received CTLA4-Ig treatment on day 8 and day 11 p.i (-TREG+CTLA4-Ig). Two treatments with GK1.5 on day 8 and day 9 p.i. were used to deplete CD4s T cells in the -CD4 group. Intravascular staining with fluorescent CD8β antibody was performed to differentiate P14 CD8 T cells in the red pulp and white pulp splenic compartments. Mice were harvested on Day 14 p.i. and analyzed with flow cytometry. A. MPEC and SLEC distribution of splenic donor antigen-specific CD8+ T cells at day 14 p.i. with LCMV_Arm_ under Treg sufficient conditions, Treg deficient conditions (with or without CTLA4-Ig supplementation), and CD4 deficient conditions. The flow plots show gating of P14 donor cells in the spleen; intravascular staining to distinguish red pulp and white pulp CD8 T cells; relative proportions of MPEC and SLEC subsets in red and while pulp splenic areas. B. GzmB expression in donor cells isolated from spleens at day 14 p.i with LCMV_Arm_ under Treg sufficient conditions, Treg deficient conditions (with or without CTLA4-Ig supplementation), and CD4 deficient conditions. The histograms and bar graphs depict the MFI values of Granzyme-B (GzmB) expression in total donors from the experimental groups. C. GzmB expression in donor cells from splenic red pulp and white pulp areas at day 14 p.i with LCMV_Arm_ under Treg sufficient conditions, Treg deficient conditions (with or without CTLAIg supplementation), and CD4 deficient conditions. The histograms and bar graphs depict the MFI values of GzmB splenic red pulp and white pulp donor cells from all the experimental groups. D. GzmB expression in donor MPECs from red and white pulp splenic compartments at day 14 p.i with LCMV_Arm_ under Treg sufficient conditions, Treg deficient conditions (with or without CTLA4-Ig supplementation), and CD4 deficient conditions. The histograms and bar graphs depict the MFI values of GzmB in donor red pulp and white pulp MPECs from the experimental groups. Data are representative of 2-5 experiments with 3-5 mice per group. Statistical significance is indicated by the following: *p ≤ 0.05, **p ≤ 0.01, ***p ≤ 0.001. p ≥ 0.05 was considered ns.

### Treg cells promote E-M transition of MPECs by suppressing Th1 responses and APC activation through the CD28-B7 axis

Given that depletion of CD4 T cells reversed the defects in E-M transition of antigen-specific CD8 T cells in the absence of Treg cells, we further characterized the nature of CD4 T cell responses in the presence or absence of Treg cells during the E-M transition phase. We found that while T_FH_ cell numbers remained unaltered, absolute numbers of CD4 Th1 cells were significantly increased in the absence of Treg cells (Fig 5A). Since Th1 cells drive the activation of APCs, which can further aggravate immune dysregulation during the E-M transition phase, we also characterized the APCs in the presence or absence of Treg cells. We found that the increase in Th1 cells was associated with a concomitant increase in the numbers of monocytic cell population, while conventional DCs (cDCs) and plasmacytoid DCs (pDCs) remained largely unaltered upon Treg cell ablation during the E-M phase (Fig 5B). Interestingly, this increase in monocytes in the absence of Treg cells was reversed by depletion of CD4 T cells or by CTLA4-Ig treatment (Fig 5B). Upon Treg ablation, the expression of CD86 and MHC-II was significantly increased in dendritic cells, and a similar trend was observed in monocytes as well, thus indicating increased activation of APCs (Fig 5C, 5D). Depletion of CD4 T cells or CTLA4-Ig treatment reversed the increased activation of APCs noted in Treg-ablated mice (Fig 5C, 5D, S5A, S5B), and also showed modest trends towards reversal of metabolic defects associated with Treg ablation, with CTLA4-Ig and CD4 T cell depletion leading to significant decrease in glucose uptake by MPECs (Fig S5C). These observations suggest that impaired E-M transition in the absence of Treg cells may be related to increased activation of effector CD8 due to increased numbers and activation status of Th cells and APCs.

**Figure 5.**
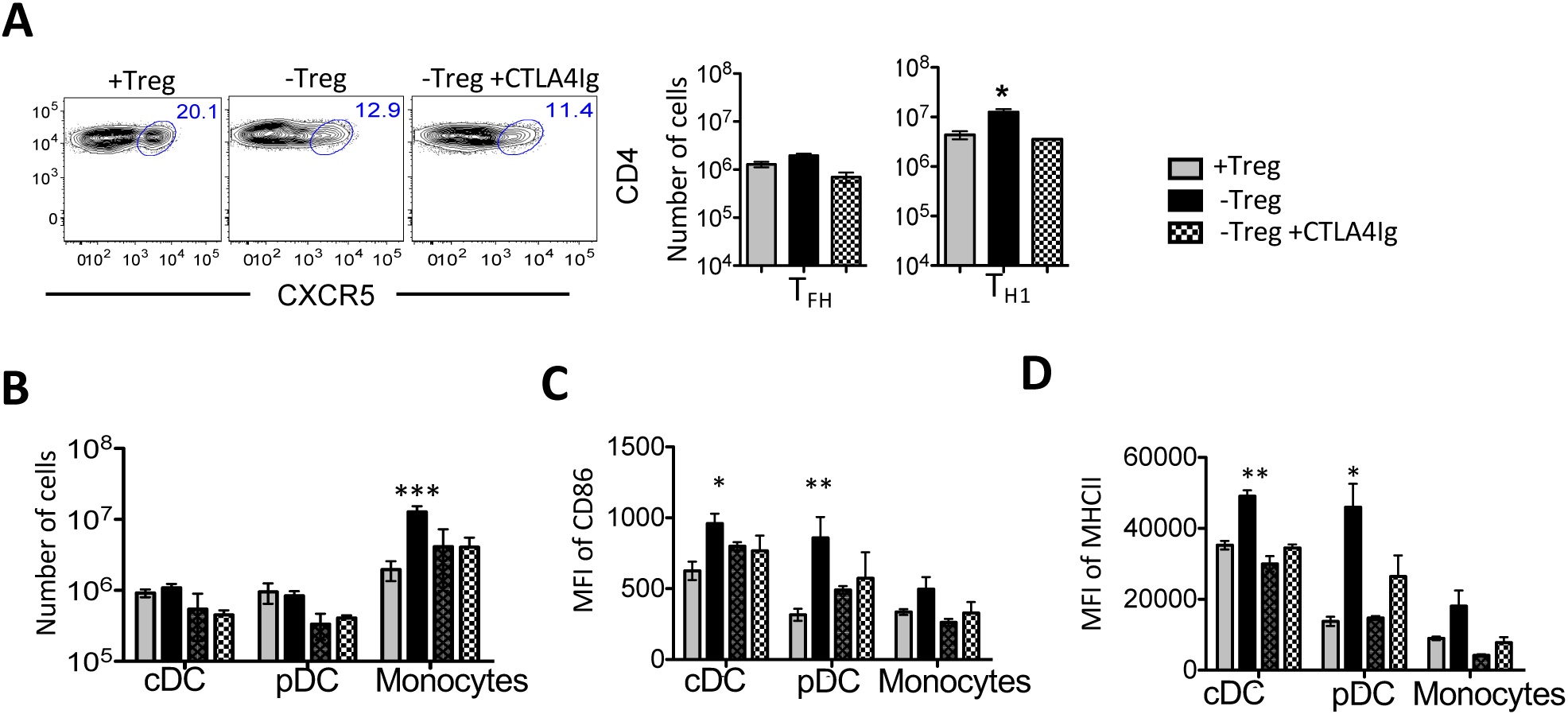
Treg cells promote effector-to-memory conversion by suppressing APC activation through effector CD4 T cells. 1×10^5^ D^b^GP33-specific P14 T cells were adoptively transferred into Foxp3^DTR^ mice, which were subsequently infected with LCMV_Arm_. Two groups of Foxp3^DTR^ mice were treated with DT on day 8 and day 11 p.i. One of these groups also received CTLA4-Ig treatment on day 8 and day 11 p.i (-TREG+CTLA4-Ig). Two treatments with GK1.5 on day 8 and day 9 p.i. were used to deplete CD4s T cells in the -CD4 group. Intravascular staining with fluorescent CD8β antibody was performed to differentiate P14 CD8 T cells in the red pulp and white pulp splenic compartments. Mice were harvested on Day 14 p.i. and analyzed with flow cytometry. A. Percentage of splenic T_FH_ cells on D14 p.i with LCMV_Arm_ under Treg sufficient and Treg deficient conditions (with or without CTLA4-Ig supplementation). Flow plots show the percentage of T_FH_ cells (CXCR5+) of the total CD4 T cell population in the various experimental groups. Bar graphs depict absolute numbers of T_FH_ and T_H1_ cells in spleen in the different experimental groups (-CD4 group not shown). B. Absolute numbers of splenic conventional dendritic cells (cDCs), plasmacytoid dendritic cells (pDCs), and monocytes on D14 p.i under Treg sufficient conditions, Treg deficient conditions (with or without CTLA4-Ig supplementation), and CD4 deficient conditions. Bar graphs depict absolute numbers pf cDCs, pDCs and monocytes in the different experimental groups C. CD86 expression in splenic cDCs, pDCs and monocytes on D14 p.i under the previously described experimental conditions. Bar graphs depict MFI values of CD86 expression in cDCs, pDCs and monocytes in the different experimental groups. D. MHCII expression in splenic cDCs, pDCs and monocytes in in spleens on D14 p.i under the previously described experimental conditions. Bar graphs depict MFI values of MHCII expression in cDCs, pDCs and monocytes in the different experimental groups.

### Treg cells promote E-M transition of antigen-specific CD8 T cells by regulating IFN-γ

IFN-γ is the primary inflammatory cytokine produced by Th1 cells, and is known to reinforce effector programs in CD8 T cells through T-be and granzyme B expression (39). Hence, we assessed the levels of IFN-γ in serum of LCMV infected mice during the E-M transition phase. We found that in absence of Treg cells, the serum concentration of IFN-γ was main-tained at significantly higher levels compared to +Treg condition (Fig 6A). Consistent with in-creased serum IFN-γ levels, IFN-γ-induced chemokines CXCL9 and CXCL10 were also in-creased in the absence of Treg cells (Fig S6A). We hypothesized that blocking IFN-γ to curb the inflammatory effect of CD4 T-cells in a Treg-depleted environment may inhibit effector pro-gram and promote memory transition of MPECs. We treated LCMV-infected mice with IFN-γ blocking antibody in the presence or absence of Treg cells during the E-M phase and analyzed the proportions and effector status of MPECs (Fig 6B). Blocking IFN- γ in the absence of Treg cells (starting from Day 8 post-infection) reversed the loss of MPECs (Fig 6B), and also prevented upregulation of effector program in the absence of Tregs as indicated by reduced levels of granzyme B expression in total donors (Fig S6B) and MPECs (Fig 6C, S6C) compared to Treg-depleted group. The granzyme B expression levels of WT and –Treg +αIFN-γ donors were not significantly different at Day 12 post-infection, while the granzyme B levels of –Treg donors were significantly elevated (Fig 6C, S6C). Together, these data demonstrate a critical role of IFN-γ in driving terminal differentiation of MPECs during the E-M phase, when Treg suppression is missing.

**Figure 6.**
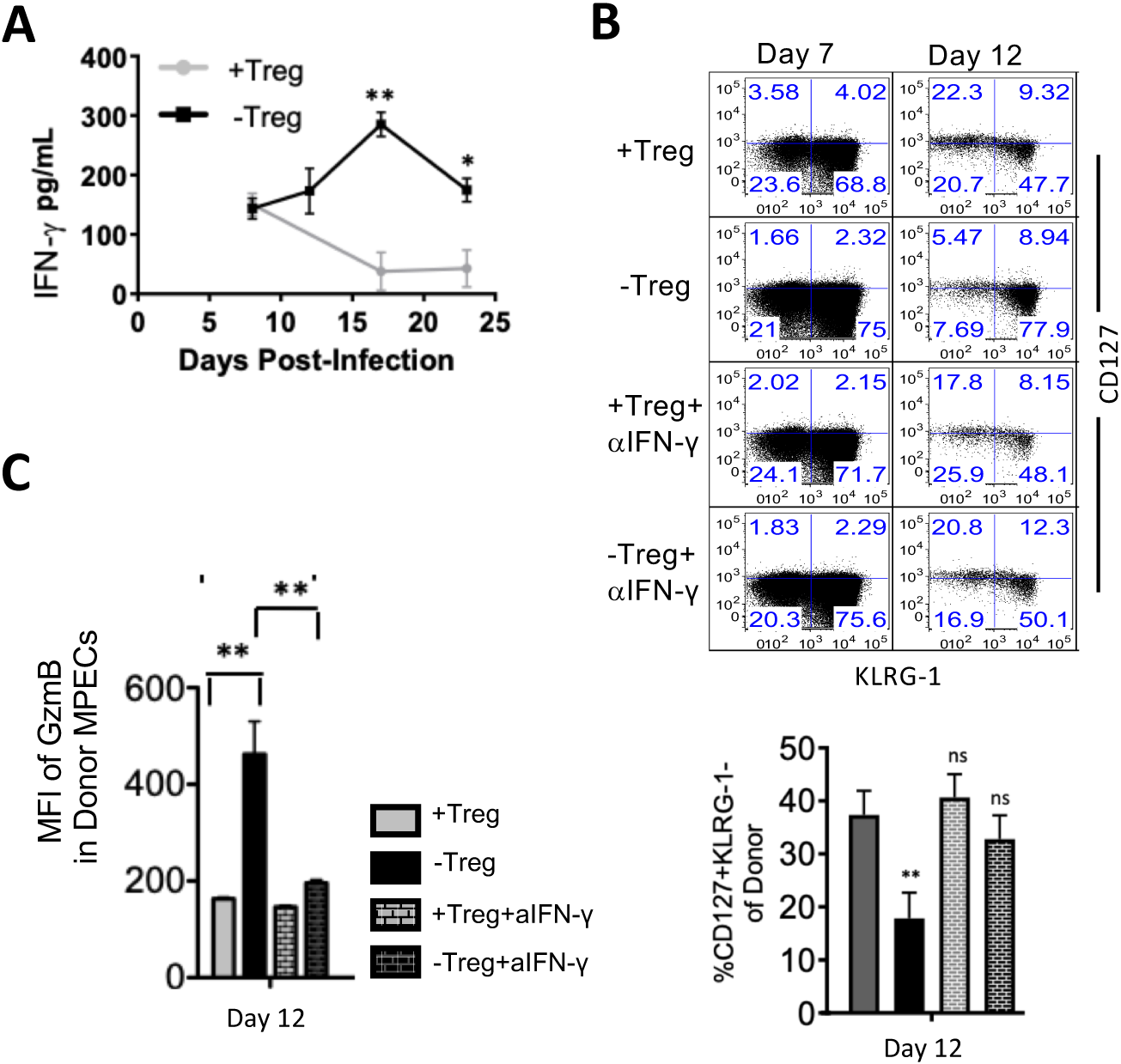
Treg cells promote effector-to-memory transition of CD8 T cells partly through suppression of IFN-γ. A. Concentration of IFN-γ in serum at indicated time points post-infection with LCMV_Arm_ in Treg sufficient and deficient conditions. 1×10^5^ DbGP33-specific P14 T cells were adoptively transferred into Foxp3^DTR^ mice which were subsequently infected with LCMV_Arm_. DT treatment was given at Day 8 post-infection to deplete Treg cells. The line graph depicts serum concentration of IFN-γ in both experimental groups at different points post-infection as assessed by Legendplex assay. B. GzmB expression in donor CD8+ PBMC MPECs on day 12 post-infection with LCMV_Arm_ under Treg sufficient and deficient conditions with or without an IFN-γ blockade. 1×10^5^ D^b^GP33-specific P14 CD8 T cells were adoptively transferred into Foxp3^DTR^ mice which were subsequently infected with LCMV_Arm_. Two groups of Foxp3^DTR^ mice were treated with DT on days 8 and 11 p.i. One of these groups was also treated with αIFN-γ on those days. Similar αIFN-γ treatment was also conducted in one +Treg group. Bar graph depicts MFI of GzmB expression in donor MPECs from the described experimental groups on day 12 post-infection. C. MPEC and SLEC distribution of donor CD8+ PBMCs on days 7 and 12 p.i. with LCMV_Arm_ under Treg sufficient and deficient conditions with or without an IFN-γ blockade. FACS plots depict the KLRG1-CD127 distribution of donor PBMC CD8 T cells at indicated time points post-infection. Bar graph shows proportions of MPECs in all experimental groups on days 7 and 12 post-infection.

## DISCUSSION

Our understanding of when and how antigen-specific CD8 T cells commit to the memory lineage is replete with key regulators such as duration and affinity of antigen stimulation, costimulatory and inhibitory receptors, cytokines, metabolites, immune cells such as CD4 T cells and Treg cells, and transcriptional and epigenetic modulators (40). However, our understanding of the mechanisms regulating the transition of memory-fated effector CD8 T cells into quiescent memory state is limited. Treg cells have been implicated in this process by several groups (18, 19, 27). In this study, we report a preferential localization of Treg cells and memory-fated MPECs in the splenic white pulp, an anatomic microniche that has been associated with the development of qualitatively superior memory cells. Global ablation of Tregs, however, led to a loss of MPEC numbers, metabolic dysregulation and an upregulation of the effector program in MPECs and SLECs alike in both the splenic red pulp and white pulp regions. This was mediated by aberrant activation of Th1 and APCs in the absence of Treg cells, and could be effectively restored by CTLA4-Ig as a surrogate functional replacement of Treg cells.

These findings support a multimodal regulatory network that promotes E-M transition through Treg-mediated suppression of effector responses either directly in CD8 T cells, and/or indirectly through suppression of Th1 responses by competing out CD28 costimulatory ligands CD80/CD86 on APCs through CTLA4. Increased IFN-γ production by effector T cells in turn may then feedforward a loop that further amplifies inflammatory immune dysregulation through increased APC activation, thereby driving terminal differentiation of MPECs in the absence of Treg cells during the E-M transition phase. Increased IFN-γ levels may also serve to further activate MPECs directly. Our studies implicate CD4 T cells as the major mediators of loss of MPEC quiescence in the absence of suppressive Treg cells. Arguably, if Tregs were exerting direct inhibitory effects on MPECs, independently of CD4 T cells, through CTLA4-mediated inhibition of CD28-CD80/CD86 costimulatory signaling in MPECs, partial MPEC defects would have been visible despite CD4 T cell depletion. Hence, the working model is that Treg cells primarily suppress Th1 cells and APCs, and in the absence of their activation an increased IFN-γ signaling, MPECs undergo quiescence.

Our observations of reduced MPEC numbers in the absence of Treg cells is consistent with previous reports of enhanced memory development in the absence of Tregs during E-M phase(18, 19, 27). Our data showing increased glucose uptake and elevated mitochondrial superoxide levels in white pulp and red pulp MPECs and SLECs in the absence of Tregs support a critical role of Treg cells in promoting E-M transition through metabolic regulation. These findings are consistent with previous reports of increased glycolysis and mitochondrial reactive oxygen species (mROS) levels in effector cells compared to memory cells (32–35). While Treg ablation was associated with similar changes in MPEC numbers, effector phenotype and metabolism in both red and white pulp splenic areas, one notable exception was mitochondrial membrane polarization. Only the red pulp MPECs and SLECs showed a change in mitochondrial polarization in the absence of Treg cells. Recent studies have shown that fused mitochondrial networks are largely seen in memory T cells, whereas effector T cells have punctate mitochondria (36–38). It is plausible that memory precursors in the white pulp are further along in fused mitochondria and are thus better able to handle the metabolic stress induced by activation in the absence of Tregs compared to the red pulp MPECs. Future studies to compare the mitochondrial dynamics of red pulp and white pulp memory precursors during this period might contain key metabolic clues to why white pulp MPECs form better memory T cells.

## ACKNOWLEDGMENTS

The authors would like to thank Shruti Bhise and Yevgeniy Yuzefpolskiy for technical assistance. Funding: This work was supported by NIH, NIAID (5R01AI132819 to SS, R21AI154363 to VK), NIA (R56AG069194 to VK), and seed funds from Seattle Children’s Research Institute (to VK and SS). Author contributions: AV, AK, YY carried out experiments, AV, AK analyzed data, prepared figures and wrote the manuscript. VK and SS conceptualized the project, designed the experiments, analyzed data, interpreted the results, prepared figures, and wrote the manuscript. Competing Interests: None. Data Availability: All data associated with this study are in the paper or supplementary materials.

## SUPPLEMENTAL FIGURES

**Supplemental Figure 1.**
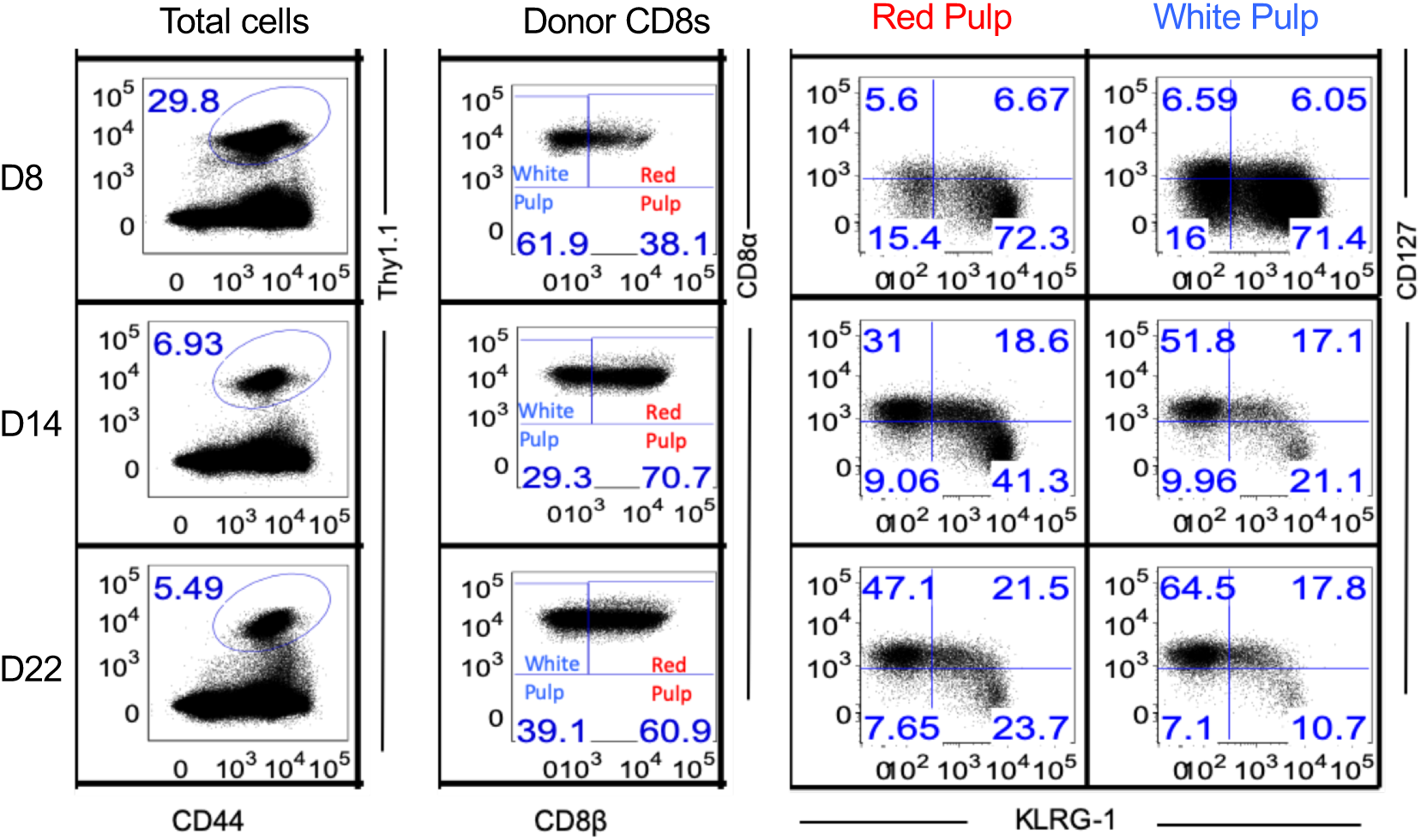
MPECs are preferentially enriched in the splenic white pulp. 1×10^5^ D^b^GP33-specific P14 CD8 T cells were adoptively transferred into C57BL/6 mice which were subsequently infected with LCMV_Arm_. Intravascular staining with fluorescent CD8β antibody was performed to differentiate P14 CD8 T cells in the red pulp and white pulp splenic compartments. Gating strategy for D^b^GP33-specific CD127^hi^KLRG-1^lo^ MPECs in splenic red pulp and white pulp compartments at days 8, 14, 22 post-infection are presented. Cell surface markers CD127 and KLRG-1 were used to assess proportions of MPEC (CD127^hi^KLRG-1^lo^) and SLEC (CD127^lo^KLRG-1^hi^) subsets as presented in the representative FACS plots.

**Supplemental Figure 2:**
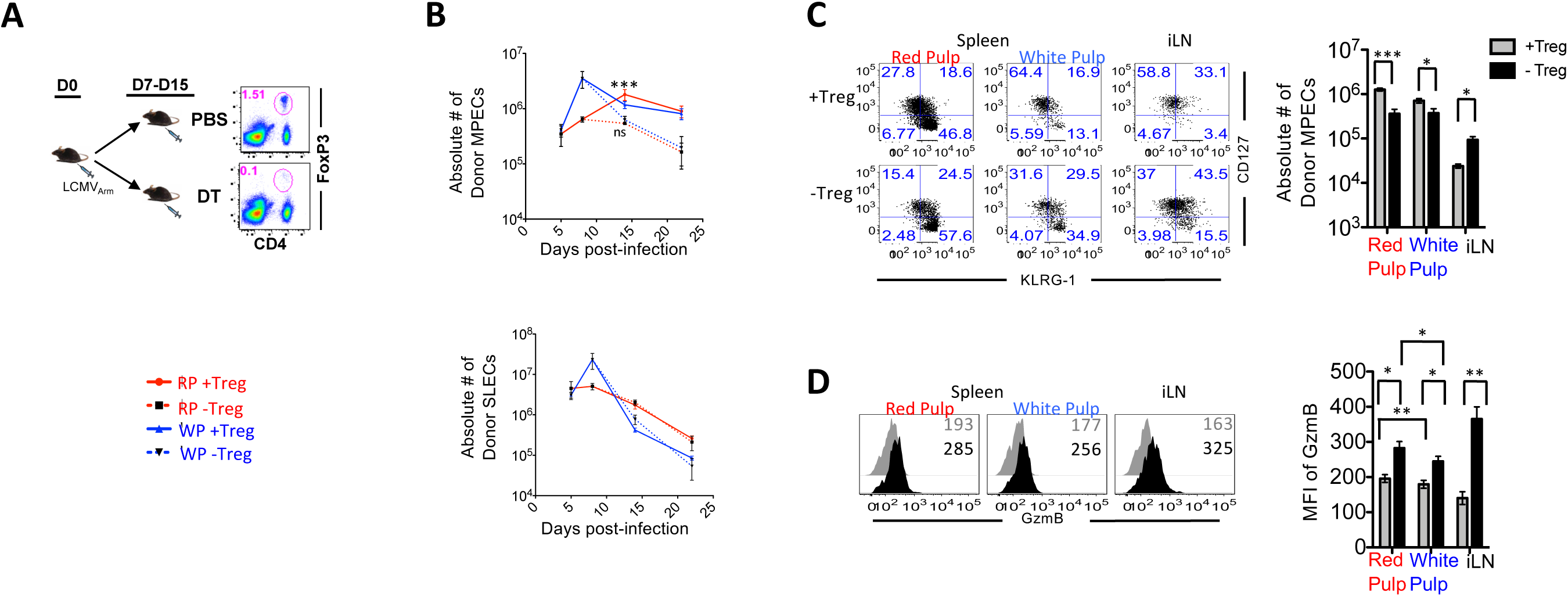
Treg cells are critical for maintenance of MPECs in the Spleen. 1×10^5^ D^b^GP33-specific P14 T cells were adoptively transferred into C57BL/6 (+Treg) and Foxp3^DTR^ (-Treg) mice which were subsequently infected with LCMV_Arm_. FoxP3^DTR^ mice were treated with Diphtheria Toxin (DT) every three days starting at day 7 or 8 to deplete Treg cells. Intravascular staining with fluorescent CD8β antibody was performed to differentiate P14 CD8 T cells in the red pulp and white pulp splenic compartments. Mice were harvested at different timepoints (D5, D8, D14, D22) and analyzed with flow cytometry. (**A)** Successful ablation of Tregs are day 15 after infection. **(B)** Absolute numbers of splenic red and white pulp MPECs and SLECs over time post-infection with LCMV_Arm_ under Treg sufficient and deficient conditions. The line graphs depict the kinetics of donor MPECs and SLECs in both splenic compartments over time. **(C)** Quantification of donor MPECs in the iLNs and splenic red and white pulp on day 14 p.i. with LCMV_Arm_ under Treg sufficient and deficient conditions. The flow plots show MPEC and SLEC distribution of the donor cells on Day 14 p.i. within the splenic compartments and the iLNs. The bar graphs depict the absolute number of donor MPECs within the different splenic compartments and iLNs in the two experimental groups. **(D)** Granzyme-B expression in donor MPECs in the red and white pulp of the spleen and the iLNs under Treg sufficient and deficient conditions. MFI values of Gzm-B expression in donor MPECs is depicted by the histograms and bar graphs.

**Supplemental Figure 3:**
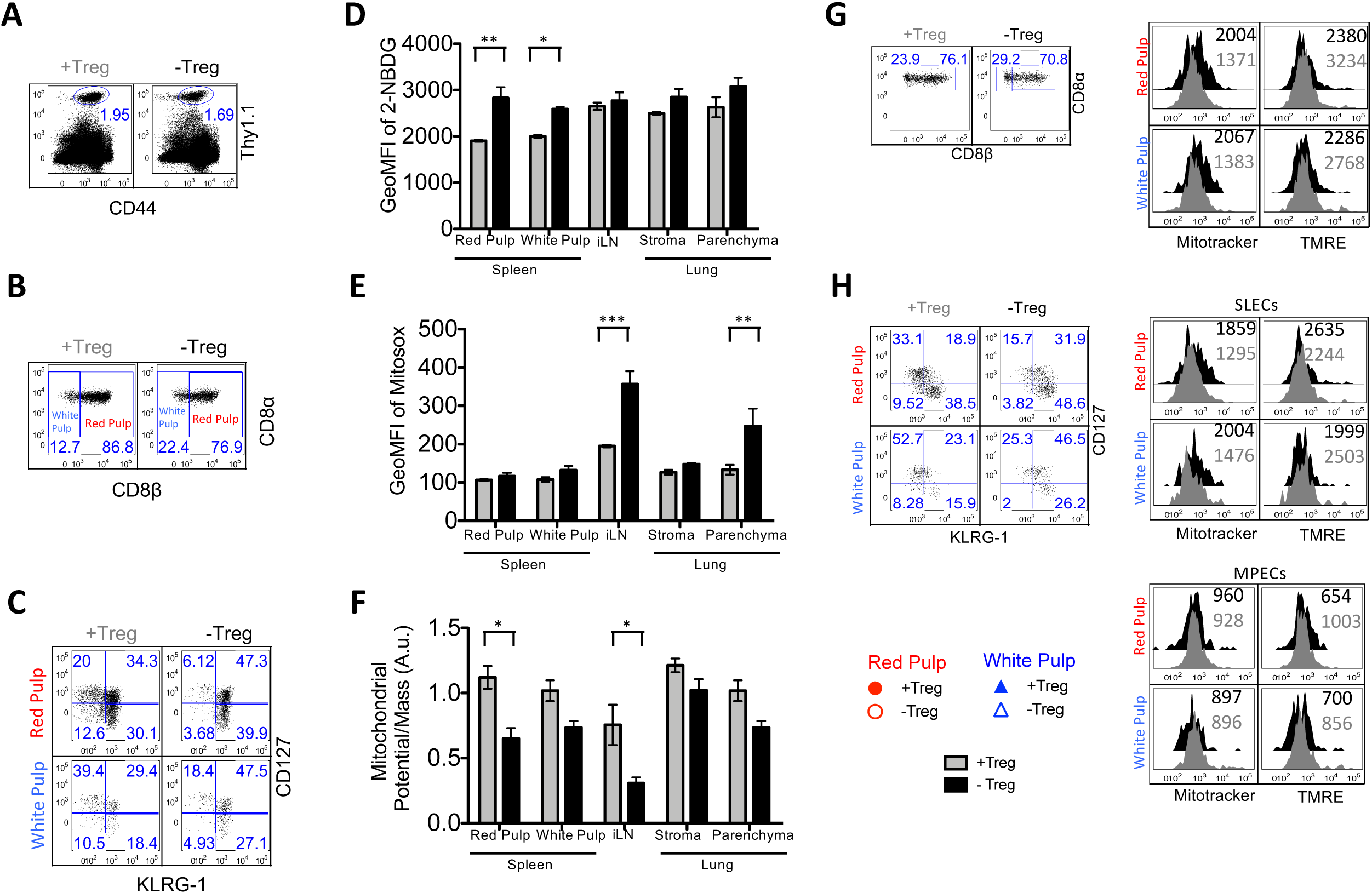
Treg cells promote healthy metabolic activity in MPECs in secondary lymphoid and nonlymphoid tissues. 1×10^5^ D^b^GP33-specific P14 T cells were adoptively transferred into C57BL/6 and FoxP3^DTR^ mice, which were subsequently infected with LCMV_Arm_. FoxP3^DTR^ mice were treated with Diphtheria Toxin (DT) every three days starting day 8 post-infection to deplete Treg cells. Intravascular staining with fluorescent CD8β antibody was performed to differentiate circulatory and tissue P14 CD8 T cells in the various harvested organs. The mice were harvested on day 14 post-infection and lymphocytes were analyzed with flow cytometry and metabolism assays. **(A)** Gating strategy to isolate splenic donor CD8+ T cells from Treg sufficient and Treg deficient conditions on day 14 post-infection for metabolic analyses. The FACS plots show donor cell gating strategy (Donors are Thy1.1+ and host T-cells are Thy1.1-). **(B)** Gating strategy to differentiate red pulp and white pulp CD8+ T cell donors from Treg sufficient and Treg deficient conditions. FACS plots display the gating strategy utilized. Red pulp donors were identified as CD8B+ and white pulp donors were CD8B-. **(C)** Gating strategy to isolate MPEC and SLEC subsets within the red and white pulp CD8+ T cell donors from Treg sufficient and Treg deficient conditions. FACS plots display the gating strategy utilized. MPECs are CD127hiKLRG-1lo and SLECs are CD127loKLRG-1hi. **(D)** Glucose uptake in donor MPECs from red pulp and white pulp of spleen, iLNs, and stroma and parenchyma of lungs at day 14 p.i. with LCMV-arm in Treg deficient and sufficient conditions. The bar graphs depict the geometric mean fluorescence intensity of NBDG in donor MPECs located in the red and white pulp of the spleen, iLNs, and the stroma and parenchyma of the lung. **(E)** Oxidative stress in donor MPECs from red pulp and white pulp of spleen, iLNs, and stroma and parenchyma of lungs at day 14 p.i. with LCMV-arm in Treg deficient and sufficient conditions. The bar graphs depict the geometric mean fluorescence intensity of Mitosox in donor MPECs located in the red and white pulp of the spleen, iLNs, and the stroma and parenchyma of the lung. **(F)** Average mitochondrial potential in donor MPECs from red pulp and white pulp of spleen, iLNs, and stroma and parenchyma of lungs at day 14 p.i. with LCMV-arm in Treg deficient and sufficient conditions. The bar graphs depict the ratio of the geometric mean fluorescence intensity of TMRE (mitochondrial potential) and Mitotracker (Mitochondrial mass) in donor MPECs. **(G)** Average mitochondrial potential in antigen specific donor CD8 T cells from splenic red pulp and white pulp compartments on day 14 p.i. with LCMV-arm in Treg deficient and sufficient conditions. The FACS plots displays the gating strategy for antigen specific red pulp and white pulp donor CD8 T cells. The histogram displays the MFI of Mitotracker and TMRE following the assay in donors from Treg sufficient and Treg deficient conditions. The scatter plot displays the MFI value of mitochondrial membrane potential per mitochondrial mass unit in donors from Treg sufficient and Treg deficient conditions. **(H)** Average mitochondrial potential in antigen specific donor MPECs and SLECs from splenic red pulp and white pulp compartments on day 14 p.i. with LCMV-arm in Treg deficient and sufficient conditions. The FACS plots displays the gating strategy for antigen specific red pulp and white pulp MPECs and SLECs. The histogram displays the MFI of Mitotracker and TMRE following the assay in donor MPECs and SLECs from Treg sufficient and Treg deficient conditions. The scatter plot displays the MFI value of mitochondrial membrane potential per mitochondrial mass unit in donor MPECs and SLECs from Treg sufficient and Treg deficient conditions. Data are representative of 2-5 experiments with 3-5 mice per group. Statistical significance is indicated by the following: *p ≤ 0.05, **p ≤ 0.01, ***p ≤ 0.001. p ≥ 0.05 was considered ns.

**Supplemental Figure 4:**
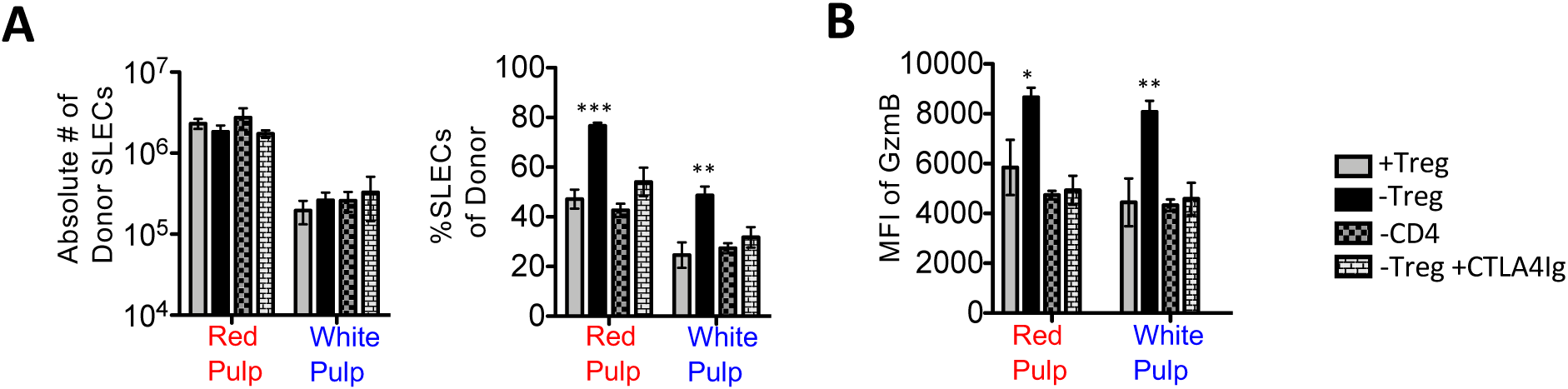
Treg Interactions with SLECs Also Occurs Through a CD4-CTLA-4 mediated mechanism. 1×10^5^ D^b^GP33-specific P14 T cells were adoptively transferred into Foxp3^DTR^ mice, which were subsequently infected with LCMV_Arm_. Two groups of Foxp3^DTR^ mice were treated with DT on day 8 and day 11 p.i. One of these groups also received CTLA4Ig treatment on day 8 and day 11 p.i (-TREG+CTLA4ig). Two treatments with GK1.5 on day 8 and day 9 p.i. were used to deplete CD4s T cells in the -CD4 group. Intravascular staining with fluorescent CD8β antibody was performed to differentiate P14 CD8 T cells in the red pulp and white pulp splenic compartments. Mice were harvested on Day 14 p.i. and analyzed with flow cytometry. **(A)** Quantification of donor SLECs from splenic red pulp and white pulp areas at day 14 p.i with LCMV_Arm_ under Treg sufficient conditions, Treg deficient conditions (with or without CTLA4Ig supplementation), and CD4 deficient conditions. The bar graphs depict both absolute number of SLECs and percent SLECs of total donor cells in the various experimental groups. **(B)** GzmB expression in donor SLECs from red and white pulp splenic compartments at day 14 p.i with LCMV_Arm_ under Treg sufficient conditions, Treg deficient conditions (with or without CTLA4Ig supplementation), and CD4 deficient conditions. The histograms and bar graphs depict the MFI values of GzmB in donor red pulp and white pulp SLECs from the experimental groups.

**Supplemental Figure 5.**
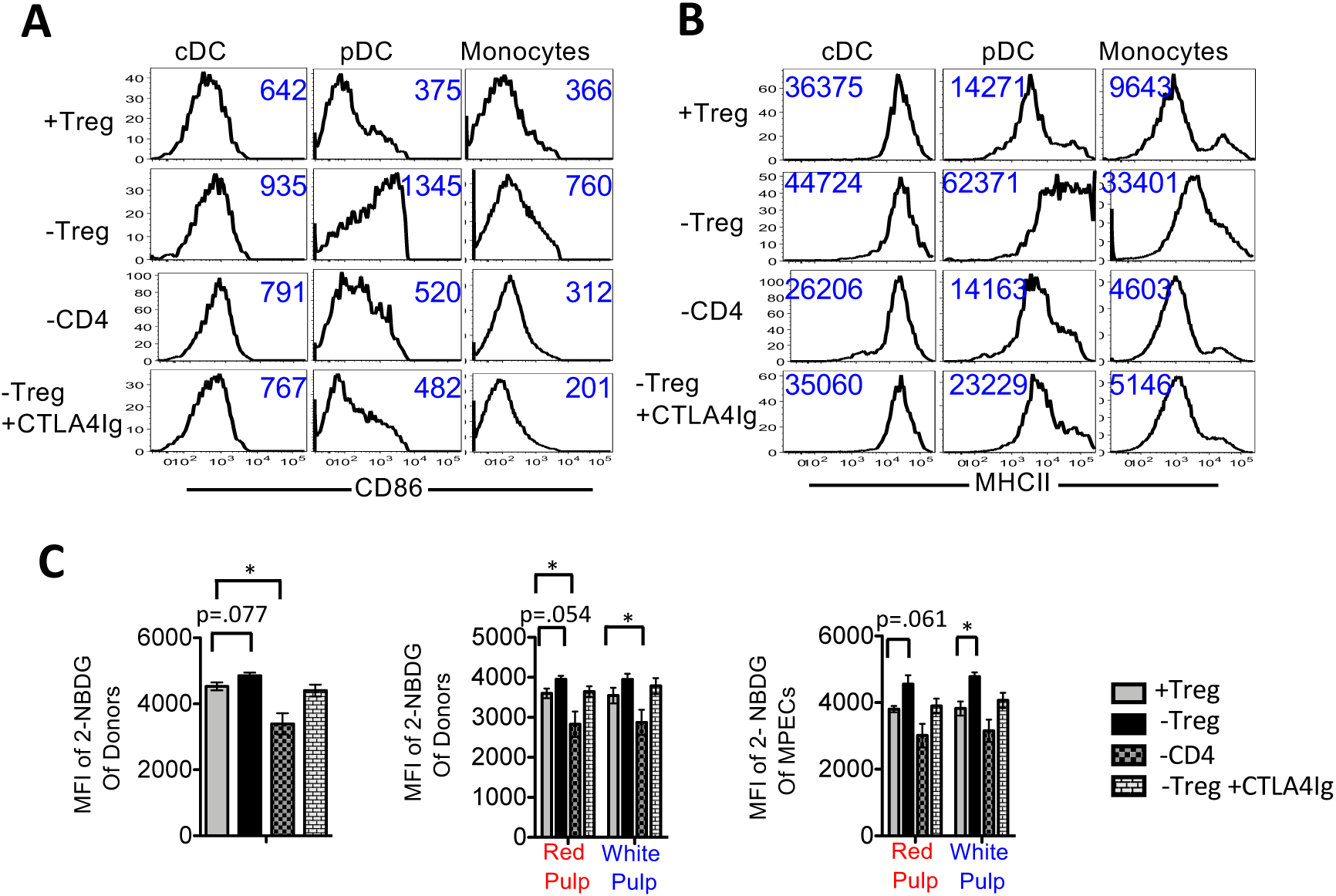
Treg cells promote effector-to-memory conversion by suppressing APC activation through effector CD4 T cells. 1×10^5^ D^b^GP33-specific P14 T cells were adoptively transferred into Foxp3^DTR^ mice, which were subsequently infected with LCMV_Arm_. Two groups of Foxp3^DTR^ mice were treated with DT on day 8 and day 11 p.i. One of these groups also received CTLA4Ig treatment on day 8 and day 11 p.i (-TREG+CTLA4ig). Two treatments with GK1.5 on day 8 and day 9 p.i. were used to deplete CD4s T cells in the -CD4 group. Intravascular staining with fluorescent CD8β antibody was performed to differentiate P14 CD8 T cells in the red pulp and white pulp splenic compartments. Mice were harvested on Day 14 p.i. and analyzed with flow cytometry. **(A)** CD86 expression in splenic cDCs, pDCs and monocytes on D14 p.i under under Treg sufficient conditions, Treg deficient conditions (with or without CTLA4Ig supplementation), and CD4 deficient conditions. Histograms depict MFI values of CD86 expression in cDCs, pDCs and monocytes in the different experimental groups. **(B)** MHCII expression in splenic cDCs, pDCs and monocytes in in spleens on D14 p.i under the previously described experimental conditions. Histograms depict MFI values of MHCII expression in cDCs, pDCs and monocytes in the different experimental groups. **(C)** Glucose uptake in donors and donor MPECs from splenic red and white pulp areas on day 14 p.i. under the previously described experimental conditions. FACS sorting was utilized to isolate donors and a glucose uptake assay was performed and analyzed by flow cytometry. The histograms show the MFI or Geometric MFI values of 2-NBDG in donor MPECs, red and white pulp donors and total donors in the indicated experimental groups.

**Supplemental Figure 6:**
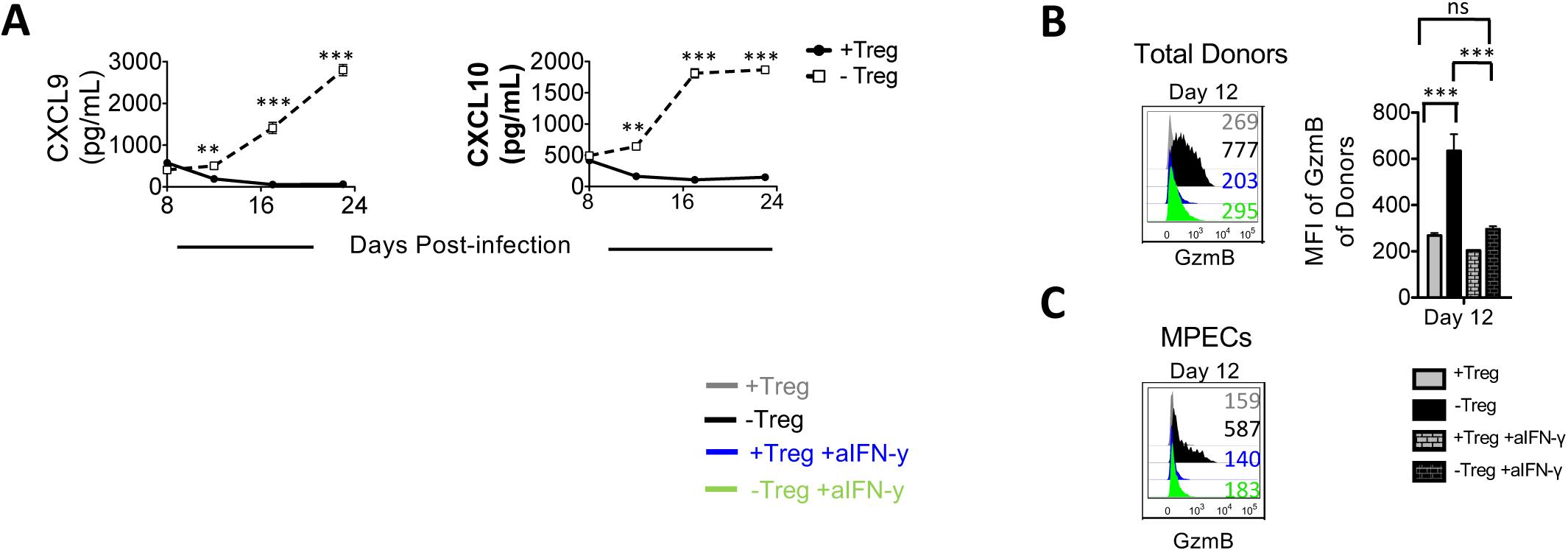
Tregs suppress production of IFN-γ inducible chemokines CXCL9/10 in a CD4-CTLA-4 dependent manner. **(A)** Serum concentrations of CXCL9 and CXCL10 after infection with LCMV_Arm_ in Treg sufficient and deficient conditions. The line graphs depict the change in concentration of CXCL9 and CXCL10 in serum over time in the presence and absence of Treg cells. **(B)** GzmB expression in donor CD8+ PBMCs on day 12 post-infection with LCMV_Arm_ under Treg sufficient and deficient conditions with or without an IFN-γ blockade. 1×10^5^ D^b^GP33-specific P14 CD8 T cells were adoptively transferred into Foxp3^DTR^ mice which were subsequently infected with LCMV_Arm_. Two groups of Foxp3^DTR^mice were treated with DT on days 8 and 11 p.i. One of these groups was also treated with aIFN-γ on those days. Similar aIFN-γ treatment was also conducted in one +Treg group. Histogram and bar graph depicts MFI of GzmB expression in donor CD8+ PMBCs from the described experimental groups on day 12 post-infection. **(C)** GzmB expression in donor CD8+ PBMC MPECs on day 12 post-infection with LCMV_Arm_ under Treg sufficient and deficient conditions with or without an IFN-γ blockade. Histogram depicts MFI of GzmB expression in donor CD8+ PMBC MPECs from the described experimental groups on day 12 post-infection.

## Notes

### Competing Interest Statement

The authors have declared no competing interest.

